# Nasal respiration is necessary for the emergence of ketamine-induced high frequency network oscillations and behavioral hyperactivity in freely moving rats

**DOI:** 10.1101/2020.07.22.213850

**Authors:** Jacek Wróbel, Władysław Średniawa, Gabriela Bernatowicz, Jaroslaw Zygierewicz, Daniel K Wójcik, Miles Adrian Whittington, Mark Jeremy Hunt

## Abstract

Changes in oscillatory activity are widely reported after subanesthetic ketamine, however their mechanisms of generation are unclear. Here, we tested the hypothesis that nasal respiration underlies the emergence of high-frequency oscillations (130-180 Hz, HFO) and behavioral activation after ketamine in freely moving rats. We found ketamine 20 mg/kg provoked “fast” theta sniffing in rodents which correlated with increased locomotor activity and HFO power in the OB. Bursts of ketamine-dependent HFO were coupled to “fast” theta frequency sniffing. Theta coupling of HFO bursts were also found in the prefrontal cortex and ventral striatum which, although of smaller amplitude, were in phase with OB activity. Haloperidol 1 mg/kg pretreatment prevented ketamine-dependent increases in fast sniffing and instead HFO coupling to slower basal respiration. Consistent with ketamine-dependent HFO being driven by nasal respiration, unilateral naris blockade led to an ipsilateral reduction in ketamine-dependent HFO power compared to the control side. Bilateral nares blockade reduced ketamine-induced hyperactivity and HFO power and frequency. In conclusion, nasal entrainment of ketamine-dependent HFO across cortical and subcortical regions at theta frequencies represents a mechanism of orchestrated neural activity across distinct brain regions. The dense divergent connectivity of the olfactory system serves to broadcast this HFO to limbic areas.

## Introduction

Bipolar depression and schizophrenia share a degree of related pathophysiology of corticolimbic circuits and genetic factors ^1,2^. Symptoms can also overlap to the extent that a schizophrenia-bipolar continuum classification has been proposed ^3^. Ketamine, at subanesthetic doses, produce a transient psychotic-like state originally reported almost sixty years ago ^4^, but paradoxically also has powerful antidepressant effects ^5^. Curiously, the acute psychosis-like effects of ketamine may be pivotal in generating its later longer-term antidepressant efficacy ^6,7^, however, ketamine’s mechanisms of action on fundamental brain network functions remain enigmatic.

Subanesthetic doses of ketamine, and related compounds, have been reported to elevate gamma oscillatory power in cortical and hippocampal regions in rodents ^8,9^, sheep ^10^, non-human primates ^11^ and humans ^12–14^. We, and others, have found subanesthetic doses of ketamine can also induce a faster rhythm (high frequency oscillations, HFO, 130-180 Hz) in diverse cortical and subcortical areas ^15–22^. There is also some evidence, from magnetencephalography, that ketamine-dependent HFO can occur in humans ^23^. Nasal respiration, has long been known to powerfully entrain brain rhythms recorded in local field potentials (LFPs) in olfactory and downstream non-olfactory areas ^24–27^. Fast oscillations (>30 Hz) also couple to this nasal respiration ^28^ and establish synchronized neuronal firing ^29^ important for neural coding of information. Slower oscillations are considered to enable integration of spatially distributed networks ^30,31^. However, it is unknown if nasal airflow drives ketamine-dependent HFO. Several rodent studies have provided descriptive or semiquantitative accounts of changes in “theta” fast sniffing after ketamine or other N-Methyl-D-aspartic acid (NMDA) receptor antagonists ^32–34^ but its relation to behavioral and fast oscillatory activity remains unknown. Given the prominent role of the OB in the generation of both nasal respiration rhythm and ketamine-dependent HFO ^35^, we tested the hypothesis that nasal airflow drives the emergence of ketamine-dependent HFO and associated behavioral activation. Since olfactory networks are closely linked with limbic areas ^36^ and since psychiatric disturbances are often associated with dysfunctional frontrostriatal connectivity ^37^ we also examined associated activity in the prefrontal cortex (PFC) and ventral striatum (VS).

## Results

### Ketamine induces fast sniffing which is reversed by haloperidol

Ketamine is known to affect NMDA receptors and dopamine systems ^38^, with dopamine associated with psychosis and to some extent depression ^2^. Rats were placed in a chamber and thermocouple signals recorded after injection of ketamine or saline. A subgroup of five rats were preinjected with haloperidol and then ketamine. Consistent with the findings of others nasal respiration typically fell into two categories, slow nasal respiration (1-3 Hz) which was typically associated with quiet waking and rest, and fast respiration (4-10 Hz) associated with exploratory activity. Rats typically switch between bouts of active sniffing lasting for several seconds to resting activity. Fig. 1 A1-3 shows example traces, and spectra shortly after injection of saline, ketamine, and haloperidol+ketamine. Stereotypic behavior, such as sniffing, is usually grouped with other repetitive behaviors ^39,40^. Using thermocouples, we were able to precisely examine nasal respiration and its relationship to behavior and oscillatory activity. After ketamine, rats displayed prolonged, almost continuous fast sniffing which lasted for around 15 min (N=8). This was visible as a change in the median sniffing frequency ~ 6 Hz compared to saline ~2 Hz. This stereotyped behavior occurred in parallel with behavioral hyperactivity. An increase in fast sniffing, a normal exploratory response, was also observed when the rat was returned to the recording chamber after saline injection, but after 1-2 min this returned to baseline values. Complete time-courses of dominant frequency and proportion of fast sniffing, along with averages of the first 15 min post ketamine are shown (p=0.001, Kruskal-Wallis, Fig. 1 B) and proportion of fast (4-10 Hz) sniffing (p<0.0001, one-way ANOVA, Fig. 1 C). In a subset of rats (N=5) we found 1.0 mg/kg haloperidol pretreatment prevented ketamine-induced increases in fast sniffing and which remained close to control values.

**Figure 1.**
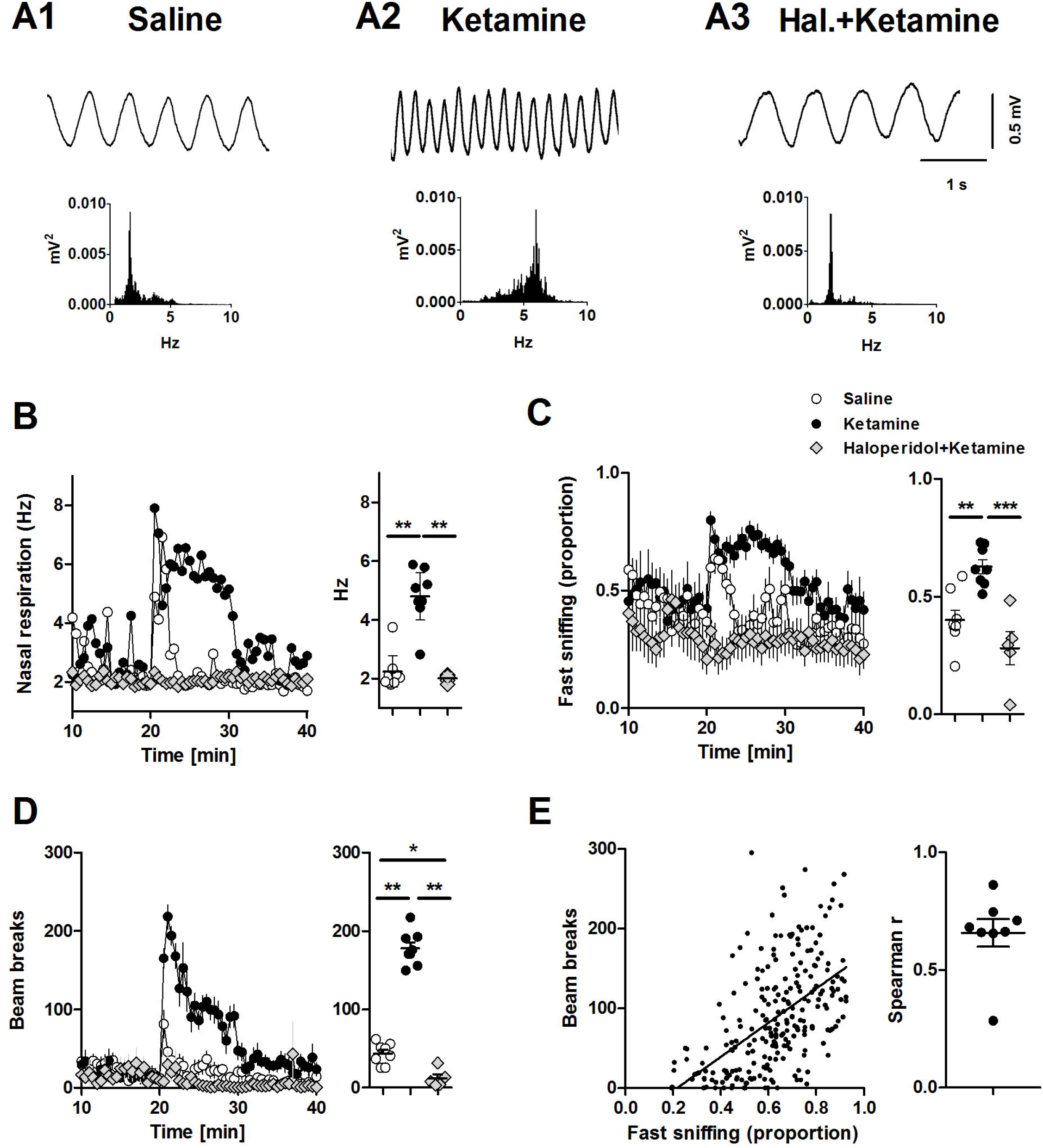
Ketamine is associated with fast sniffing which is reversed by haloperidol. A1-3, example traces of nasal respiration shortly after injection of saline, ketamine or haloperidol+ketamine. Power spectra calculate for 60 s are also shown. B, Time-courses showing the median dominant sniffing frequency after injection of saline, ketamine (N=8 rats) and haloperidol+ketamine (subset N=5 rats). Adjacent, shows the dominant sniffing frequency for each rat calculated for the first 15 minutes after injection (p=0.001, Kruskal-Wallis, Dunn’s post hoc). C, Time-courses showing the proportion of fast sniffing after injection. Adjacent shows proportion of fast sniffing (4-10 Hz) for each rat calculated for the first 15 minutes after injection (p<0.0001, one-way ANOVA, Bonferroni post hoc). D, Time-course showing beam break activity after injection. Adjacent shows mean beam breaks for each rats calculated for the first 15 minutes after injection (p<0.0001, one-way ANOVA, Bonferroni post hoc). E, Correlation between the proportion of fast sniffing and beam break activity for the first 15 min post injection for all rats. Linear correlation slope differed from zero (p<0.0001, one-way ANOVA). Adjacent shows the Spearman’s rank correlation score which was significant for 7 of 8 rats. *p<0.05, **p<0.01, ***p<0.001.

Ketamine behavioral hyperactivity shared a similar time course with fast sniffing and was also reduced by 1.0 mg/kg haloperidol pretreatment (Fig. 1 D). Pooled data from all rats for first 15 min post ketamine revealed a positive correlation between beam breaks and the proportion of fast sniffing (Spearman r=0.7366; p<0.0001). Analysis of individual rats revealed correlations were significant in 7/8 rats (p<0.001). Prolonged fast sniffing was present in one rat in which no correlation was found (Fig. 1 E).

### Ketamine increases HFO power which correlated with fast sniffing

We next examined oscillatory activity within the LFP recordings from the OB (N=8) in more detail. In control conditions, HFO (130-180 Hz) are often barely visible in spectra (Fig. 2 A1). By contrast, gamma oscillations are prominent features of LFP’s recorded from the OB and divided into low- (30-70 Hz) and high-gamma bands (70-100 Hz) ^41^. Consistent with previous reports ketamine produced an almost immediate, selective increase in HFO power lasting around 15 minute before returning to baseline (Fig. 2 A2) ^15,18,19,22,35^. Time courses of the effect of intraperitoneal injection of ketamine on low-gamma and high-gamma, and HFO are shown in Fig. 2 B1. Analysis of the mean 15 min before and immediately after ketamine revealed a significant increase in HFO power (p=0.0078 paired-test). Changes in low- and high gamma did not reach significance (p=0.098, p=0.20, respectively, paired t-test, Fig. 2 B2). Saline injection did not influence gamma or HFO power. Given the striking similarities of the time-courses of HFO, fast sniffing and locomotion we examined if there was a relationship between these measures. Pooled analysis from all rats of the first 15 min post ketamine revealed a significant correlation (Spearman) between HFO power and proportion of fast sniffing (r=0.3; p<0.0001) and a stronger correlation for HFO and beam breaks (r=0.6; p<0.0001). Analyses from individual rats revealed this correlation was significant for HFO power/fast sniffing in 7/8 rat and HFO power/beam breaks in 8/8 rats (individual R values are shown in Fig. 2 C).

**Figure 2.**
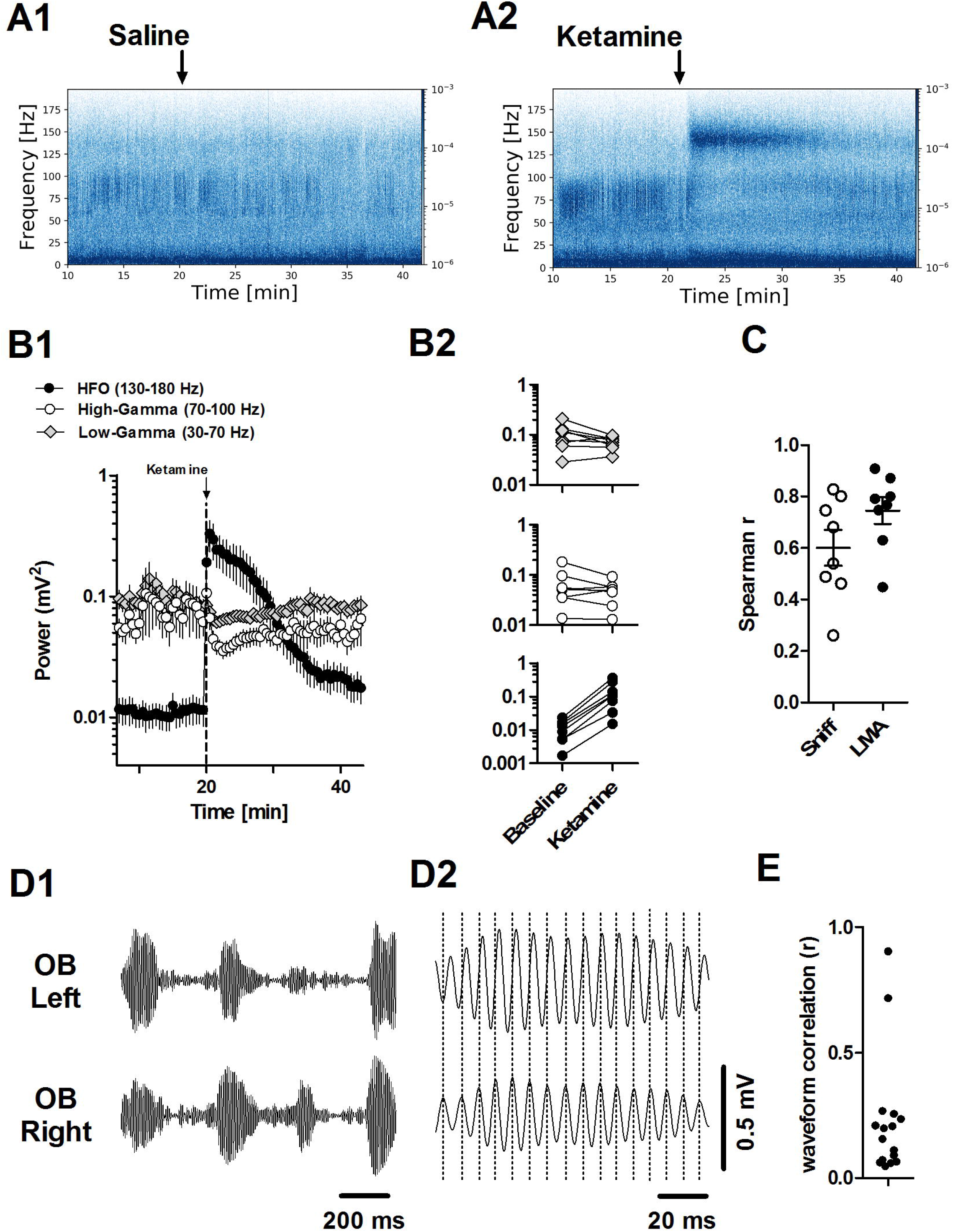
Ketamine differentially affects gamma and HFO power. A1-2, Example spectrograms showing the effect of vehicle and ketamine injection on LFP oscillations recorded in the OB. B1, Time-courses showing the effect of ketamine injection on the power of low (30-70 Hz)- and high (70-100 Hz) gamma and HFO (130-180 Hz). B2, Plots for individual rats showing the mean power at baseline and mean power post injection of ketamine (15 min blocks, N=8 rats as in Fig. 1). C, Scatter plot showing the correlation between HFO power and proportion of fast sniffing (white circles) and beam breaks (black circles). Spearman r values are shown for each rat. D1, 130-180 bandpass-filtered LFPs from the left and right OB. Note, bursts of HFO, after ketamine, are largely coherent on both sides, however occasional bursts may occur independently on one side. D2, expansion of a single burst showing that although bursts are often coherent they are not synchronized. E, 130-180 Hz waveform correlation values from the rats for left vs. right sides, median and range is shown.

Bursts of ketamine-HFO were largely coherent on left and right sides, however, occasional asymmetric bursts could be observed (Fig. 2 D1). Closer inspection revealed that although bursts were largely coincident they were not synchronous with troughs rarely in phase between sides (Fig. 2 D2). Waveform correlation for left and right OB revealed weak correlation scores for all rats (Fig. 2 E) indicating independent HFO generators each driven by airflow through each nostril separately.

### Ketamine-induced HFO is coupled to nasal respiration

Considering ketamine influences nasal respiration and that nasal respiration can entrain oscillations in the OB we investigated this relationship further (analyses based on thermocouple/LFP data; N=8). Example simultaneous thermocouple recordings, LFPs (1-10 Hz, 70-100 Hz, and 130-180 Hz) and average spectra are shown in Fig. 3 A, B. At baseline we observed significant coupling for 70-100 Hz oscillations to the thermocouple signal and local OB oscillations. The driving frequency was 5-8 Hz, which corresponded to fast sniffing (Fig. 3 B1). Immediately after injection of ketamine we observed significant coupling to the HFO band at the same driving frequency (Fig. 3 B2). Notably, bursts of HFO and gamma occurred at different phases of the respiratory rhythm, with HFO associated towards the peak and gamma on the ascending phase (see polar plots and Supplementary Fig.1).

**Figure 3.**
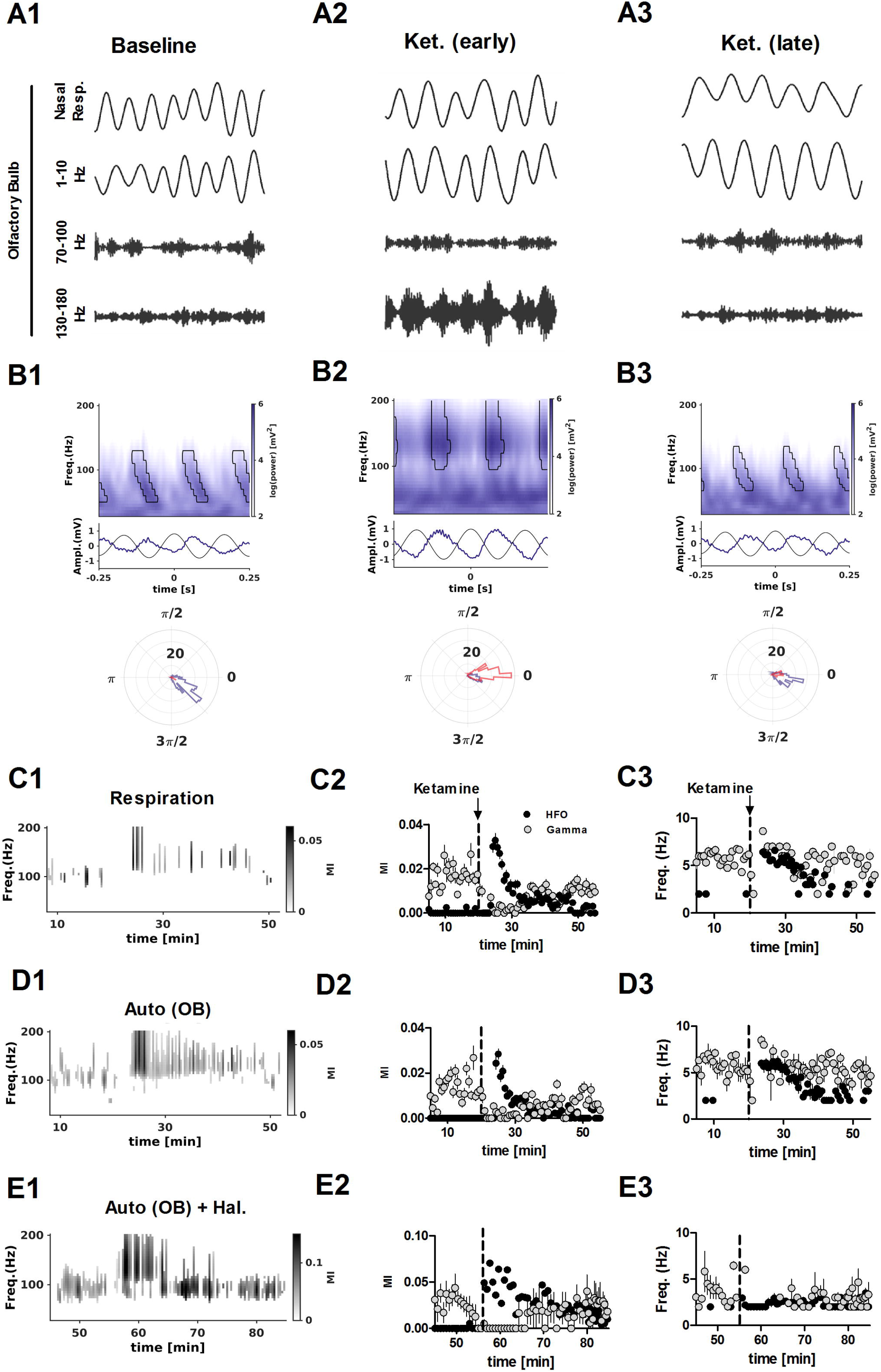
Nasal respiration entrains ketamine HFO in the OB. A1-3, Example of simultaneously-recorded nasal respiration and LFP oscillatory activity at baseline and after injection of ketamine. B1-3, Frequency spectra showing the relationship between dominant oscillatory activity in the OB and both thermocouple (black waveform) and local 1-10 Hz activity (blue waveform). Adjacent polar plots show the phase of high-gamma (blue) and HFO (red) with respect to local OB oscillations. Note high-gamma tends to occur on the ascending phase, while HFO is associated with the peak. C1, Example time course showing strength of only significant (p<0.05 with False Discovery Rate correction) cross frequency coupling of fast LFP rhythms to nasal respiration C2, Mean and SEM of the modulation index (MI) for all rats and C3, corresponding driving frequency (N=8 rats as in Fig. 1). Note, dominant gamma coupling (grey) and negligible HFO coupling (black) at baseline, however both are driven by a 6 Hz “theta” frequency. D1-3, same as C except coupling was evaluated in relation to local 1-10 Hz activity from the same channel. E1-3, same as D except the rat received an initial injection of haloperidol 1 mg/kg (N=5 rats).

Time-courses showing significant coupling of fast rhythms to 1-10 Hz activity from an example rat are shown, along with mean modulation index (MI) and mean driving frequency for all rats in Fig. 3 C, D, E. After injection of ketamine, respiratory and auto coupling (coupling within the same LFP signal) to high-gamma diminished and instead strong coupling to HFO was observed. As the ketamine effect wore off (around 20 min.) coupling returned to pre-ketamine levels (Fig. 3 C1-3, D1-3). In the rats pretreated with haloperidol, injection of ketamine was also associated with HFO coupling but at a slower driving frequency, around 2 Hz and this coupling was stronger than ketamine alone (mean of first 15 min. p<0.05, paired t-test, Fig. 3 E1-3).

### Ketamine-dependent HFO bursts are synchronous across ipsilateral brain regions and modulated by theta frequencies

We examined the relationship between ketamine-dependent HFO in the OB and VS and PFC, since these regions have been shown previously to display increases in HFO power after ketamine ^15,19^. In a separate series of rats (N=6) implanted with electrodes in the OB, VS and PFC injection of ketamine was associated with bursts of HFO that shared similar temporal profiles (Fig. 4 A1). Intraburst analyses revealed the bursts were almost synchronous on a cycle by cycle basis (Fig. 4 A2). Waveform correlations revealed almost zero phases lag between OB and PFC and a consistent but small offset between OB and VS although coupling was stronger in VS (mean of first 10 min post ketamine, Fig. 4 A3).

**Figure 4.**
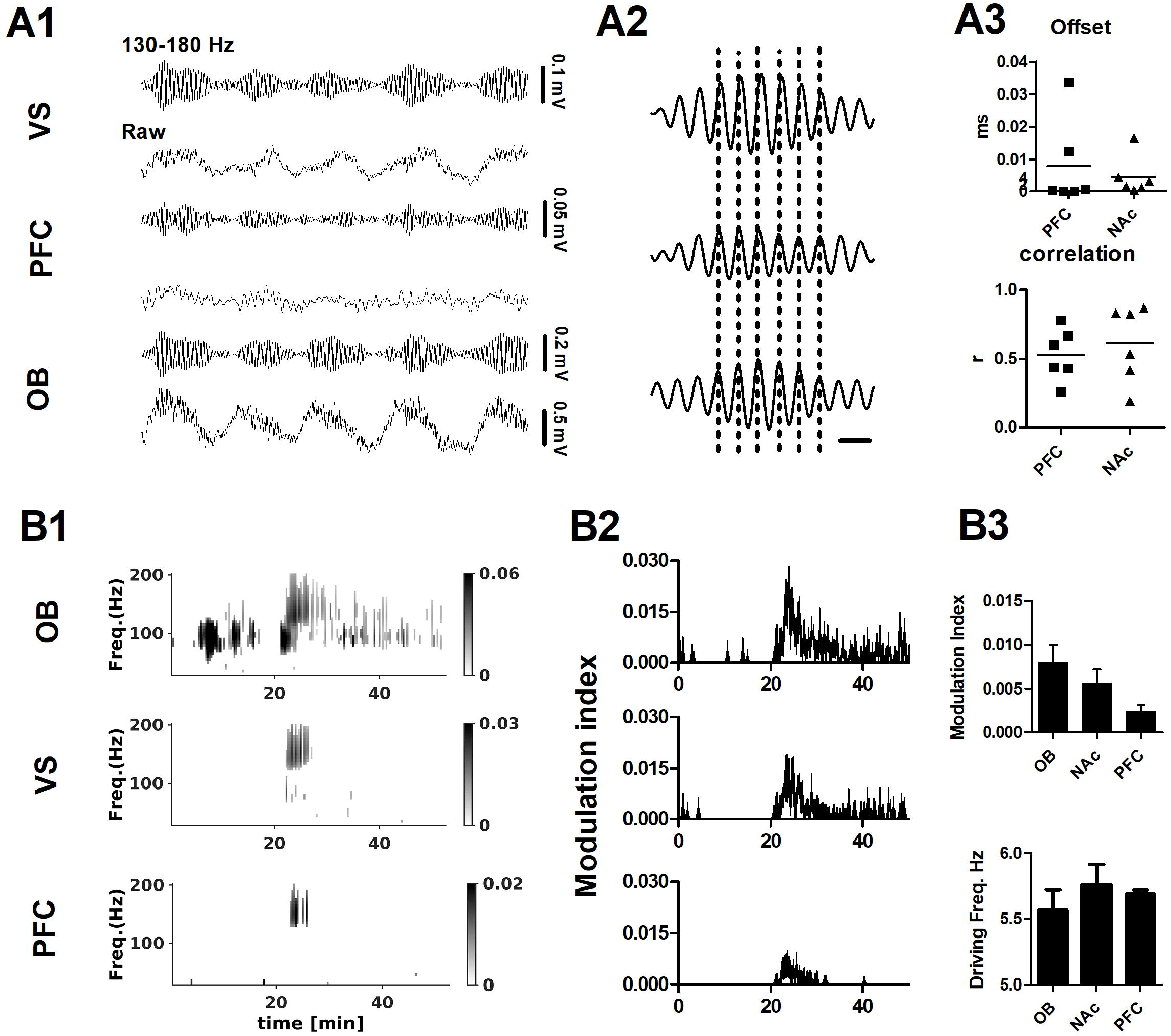
Ketamine-dependent HFO in multistrutures are dependent on nasal airflow. A1, Example raw and 130-180 Hz bandpass-filtered waveforms (1 second) from the OB, PFC and VS after injection of ketamine (N=6 rats). An expanded HFO burst is shown in A2 showing close to zero-phase lag between the OB and PFC and VS, bar indicates 5 ms. A3, mean offset between cycles of HFO in the PFC/VS with respect to OB (peak-to-peak differences for the first 10 min post injection). Waveform correlations for the 130-180 Hz signals in the OB with respect to PFC/VS (correlation for first 10 min post injections). B1, Example time course showing strength of only significant (p<0.05 with False Discovery Rate correction) coupling for local 1-10 Hz rhythms and fast oscillations for the OB, PFC and VS. B2, Modulation index (MI) for mean and SEM for all rats N=6. B3, Mean MI, and driving frequency (first 10 min post ketamine).

Olfactory networks are closely linked with limbic areas ^36^. This suggests nasal breathing may influence oscillatory activity in downstream limbic areas. After injection of ketamine significant theta-HFO coupling was observed in all structures (see Fig. 4 B1 for an example and mean MI time courses in Fig. 4 B2). Two-way ANOVA (group×time F_314, 2370_=4.68, p<0.0001) revealed coupling was significant stronger in the OB vs. VS and PFC (all p<0.001). Mean MI and driving frequency ~5.5 Hz for first 10 min are shown in Fig. 4 B3.

### Unilateral naris blockade reduces ketamine-induced HFO in cortical and subcortical areas

Since nasal respiration and ketamine-HFO are critically linked we tested the hypothesis that unilateral naris occlusion reduces ketamine-HFO power. Rats with electrodes implanted bilaterally in the OB (N=8) and VS, PFC (N=7) received unilateral nares blockade. Baseline respiratory rhythm was visible in OB recordings however, although it was not always as pronounced in the raw LFP in VS and PFC channels. Unilateral naris occlusion was associated with almost immediate desynchronization of the raw LFPs on the ipsilateral side (Fig. 5 A1, B1, C1), as reported by others ^42^. This served as a reliable marker of naris blockade. Unilateral naris occlusion was associated with diminished HFO power after ketamine on the ipsilateral side, whilst on the control side HFO remained large time×group OB (F_59,590_=2.52, p<0.0001), VS (F_59,708_=2.52, p<0.0001), PFC (F_59,708_=2.52, p<0.0001). Inset histograms shows the HFO modulation index for the first 15 min post ketamine injection and shows that this was also significantly reduced on the ipsilateral versus contralateral side (Fig. 5 A2, B2, C2).

**Figure 5.**
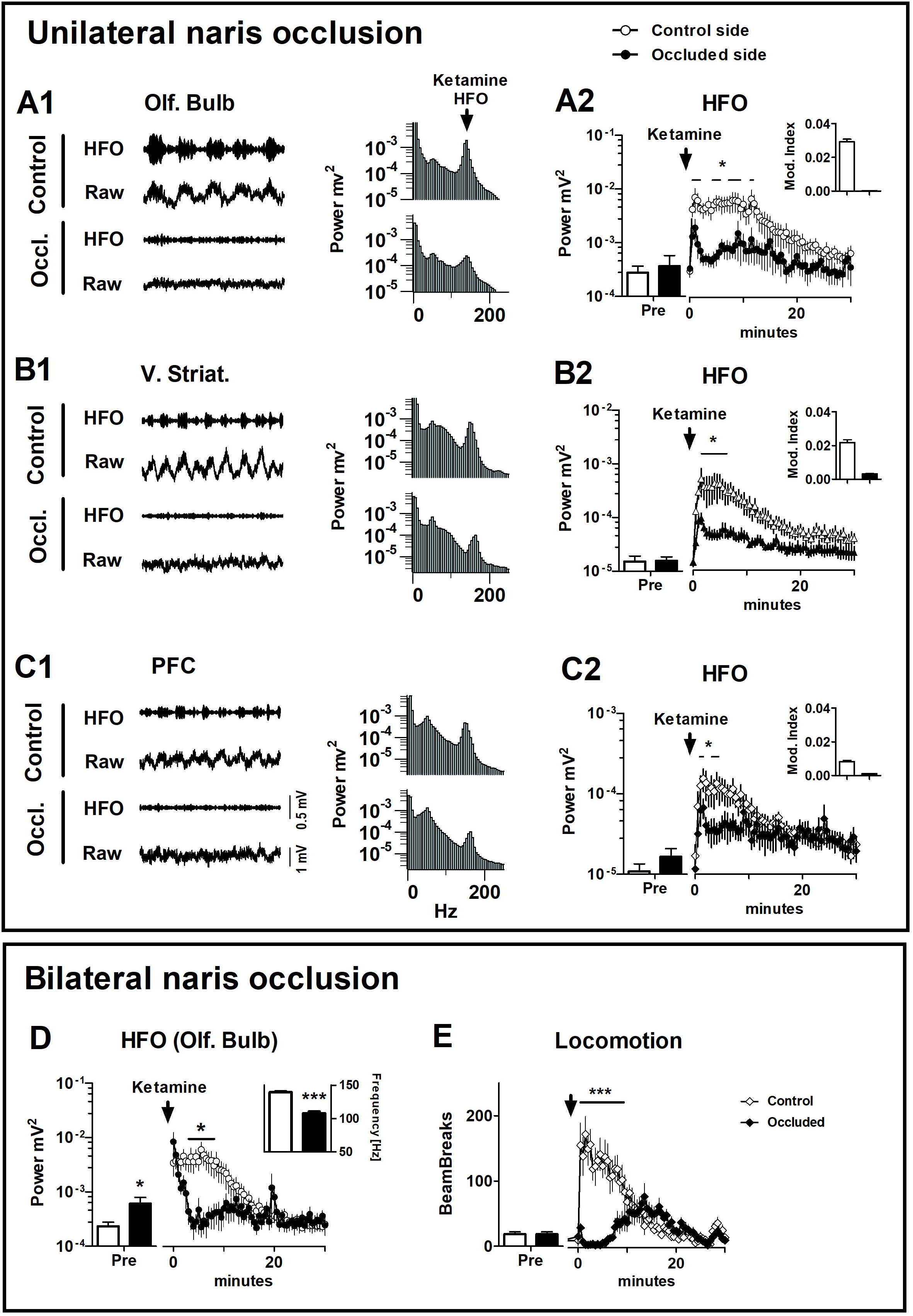
Effect of naris blockade on ketamine-induced HFO and hyperactivity. A1, B1, C1, Raw LFPs and 130-180 Hz band-pass filtered waveforms post ketamine recorded bilaterally from the olfactory bulb (N=8), ventral striatum and prefrontal cortex (N=7). In each case the occluded (ipsilateral) and control (contralateral) waveforms are shown. Corresponding 60 sec power spectra following ketamine injection are also shown. A2, B2, C2, Time courses for all rats showing unilateral naris blockade reduced the power of HFO in the olfactory bulb, ventral striatum and PFC. Note Y-axes are sequentially an order of magnitude smaller. Inserts for each time course show the modulation index for occluded versus control sides for the first 14 min. post injection of ketamine. D, time courses of ketamine-induced increases in HFO power. Bilateral naris occlusion was associated with a significant reduction in HFO power compared to the control (non-occluded) state (p<0.05, 2 way-ANOVA, Bonferroni post hoc). Although the power of spontaneous HFO was equivalent in control and occluded groups, post occlusion unexpectedly we observed a small but significant increase in the power of HFO. Inset, shows the HFO frequency (initial 5 min period post ketamine) which was significantly reduced in the naris occluded group. E, time courses of beam breaks after ketamine in the for bilateral naris occluded state and for control conditions. Bilateral naris occlusion was associated with a significant reduction in beam breaks which was also notably delayed occurring several minutes after ketamine injection. Bilateral naris occlusion did not influence beam break activity prior to ketamine injection.

### Bilateral naris blockade reduces ketamine-induced hyperlocomotion and produces complex effects on HFO

Respiratory rhythms, which were clearly visible in the OB LFPs at baseline were almost completely absent in both OB’s after occlusion. Interestingly, we did observe a small increase in the power of a fast-oscillatory rhythm, around 100-120 Hz. In bilateral nares-occluded rats, ketamine injection was associated with an immediate increase in the power of HFO, lasting 1-2 minutes followed by a return to baseline levels. In controls, ketamine-induced increases in the power of HFO lasted for the standard length of time, 10-12 minutes. Repeated measures two-way ANOVA revealed a significant time × group interaction (F_79,1106_=3.79, p<0.0001). Bonferroni post hoc analyses revealed a significant reduction of HFO power in occluded rats for the first 3.5-9 min post ketamine injection (p<0.05; Fig. 5 D). Unexpectedly, dominant frequency after ketamine was markedly reduced in occluded rats (107.9±2.5 Hz, mean of the first 5 min) after injection compared to 139.3±1.5 Hz in controls (p<0.001, paired t-test, Fig. 5 D insert).

Considering that the power of ketamine-induced HFO and locomotion correlate positively, and that unilateral nares blockade can reduce HFO, we speculated that bilateral nares blockade would impact ketamine-induced locomotion. In rodents, bilateral nares blockade was associated with clear mouth breathing, which was present throughout the course of the recording. In nares-occluded rats, ketamine-induced behavior was qualitatively and quantitatively different compared to controls. Rather than the typical circling behavior, associated with ketamine in controls, we observed an initial ataxic response where the rat would lose balance, ‘wobble’, and slowly move its head from side to side. This lasted for around 5 minutes and was followed by mild locomotor excitation. Compared to controls, this hyperlocomotor response was delayed in occluded rats and significantly less intense. Repeated measures two-way ANOVA revealed a significant effect of group (F1,79=12.12, p=0.0037) and time×group interaction (F79,1106=11.78, p<0.0001). Bonferroni post hoc analyses revealed a significant difference for the first 10 min post ketamine injection (p<0.001, Fig. 5 E). Notably, we also examined ketamine-induced locomotion for unilateral naris occlusion and there were no significant differences compared to non-occluded controls (F1,79=0.01, P=0.9, Supplementary Fig. 2).

## Discussion

In control conditions both low- and high-gamma were present in OB LFPs, but clear theta coupling was found only for the high gamma band (around 80 Hz). A finding consistent with observations of others who have associated this coupling with exploration and sensori-motor processing ^41,43^. Low-gamma oscillations in the OB have been associated with grooming and quiet waking without significant coupling ^41^. Ketamine increased the power of the faster HFO band which was coupled to theta rhythms. HFO-theta coupling was strongest in the OB, but also present in the VS and PFC. Thus, with respect to fast oscillations, ketamine could be said to have two effects. Firstly, the natural theta-gamma coupling, considered important for neural coding ^44^ would be derailed. Secondly, the emergence of aberrant HFO which correlated with two key features of the ketamine model, stereotypic sniffing and behavioral hyperactivity. Notably, ketamine-HFO interact with atypical antipsychotics leading to a reduction in frequency, of around 80 Hz ^45^, which may reflect the system trying to reset back to baseline gamma levels.

Abnormal oscillatory activity have been associated with psychiatric diseases, perhaps most notably schizophrenia. Ketamine is reported widely to affect brain rhythms in experimental animals and humans. Gamma rhythms have received a lot of attention due to their role in higher order functioning and increases gamma band activity speculated to represent elevated background electrophysiological noise with translational value for models of schizophrenia 46. Jones and colleagues have shown that clozapine, olanzapine, and haloperidol reduce keta-mine-dependent increases in ECoG gamma power ^47,48^. The HFO band also interacts with antipsychotics, chiefly second generation drugs, but in a different way, reductions in frequency rather than power occur. There are important differences between gamma and HFO, for example gamma can be recorded in a number of paradigms in the drug free state, and has widely reported associations with higher cognitive functions. In contrast, HFO is extremely weak at baseline and although increased during quiet waking and REM sleep clear functional links remain unknown. A different picture emerges after NMDA receptor administration, where increases in HFO power tend to be the dominant fast oscillatory change in many brain regions, of awake rodents, and are visible as a clear peak in the power spectra. It is worth pointing out that ketamine-dependent HFO, although of similar frequency, occur independently of hippocampal ripples ^16^ appear to involve larger scale ventral olfactory/limbic networks. With respect to gamma, in experimental rodents, NMDA receptor antagonists increase gamma power in most brain regions, which tend to be broadband and in many cases smaller than those reported for HFO. However, reductions in gamma power have also been observed after ketamine in the nucleus accumbens (bipolar LFPs) ^16^, after microinfusion of MK801 to the OB ^49^, and also in the present study, indicating that ketamine produces regionally-specific effects on different oscillatory networks.

The mechanisms underlying presence of coherent ketamine-dependent HFO across different brain regions are unclear. This rhythm might be attributed to local HFO generators e.g. in the VS ^50^ dependent on OB input ^35^. Indeed, mitral tufted cells, the main projection neurons from the OB, project to diverse regions and can impose oscillatory activity in their targets (e.g. gamma in the piriform cortex ^51^). Volume-conducted signals can be detected many millimeters from their site of generation ^52^ and may explain the presence of fast rhythms in VS ^53^. Since ipsilaterally-recorded ketamine-dependent HFO were almost perfectly in phase and of considerably smaller amplitude in the VS/PFC (compared to OB), it also possible part of this rhythm arises from volume-conducted currents.

Fast oscillations are ubiquitous in sensory networks and can synchronize spike discharge of neuronal ensembles ^54^. Coupling of faster with slower oscillations is considered to enable communication between distant brain areas. In this study we found synchronous bursts of ketamine-dependent HFO in the OB, PFC and VS coupled to theta-driven respiration. Coupling of fast brain rhythms to theta frequencies has been linked functionally to working memory ^31^, visual ^55^, and olfactory processing ^56,57^. Ketamine, which enables local networks to generate HFO more easily, combined with nasal respiration represents a mechanism of synchronization across brain regions, with each respiratory cycle (driven by a wave of stimuli hitting the olfactory system).

Our study shows that ketamine induced a distinct breathing pattern in rats. Unlike natural sniffing, ketamine-induced sniffing was continuous, for almost the complete duration of ketamine’s action and appeared stereotyped and purposeless. This finding is consistent with the general increases in stereotypy reported after ketamine and related NMDA receptor antagonists ^58^. Increases in nasal respiration are generally linked to locomotor activity and have been associated with exploration ^59^, reward expectancy ^60^ and increased metabolic demand ^61^. In line with this, we observed ketamine-dependent increases in sniffing and locomotion were correlated both with each other and the increases in HFO power. Stereotypies and hyperlocomotion are classical dopamimetic behaviors ^62^ and the dopamine agonist apomorphine has long been known to produce stereotypic sniffing in rodents ^63^. By contrast, HFO power is not markedly affected by dopamimetics ^15^, including apomorphine (unpublished) or by dopamine receptor blockade ^45^, indicating NMDA receptor blockade is the key factor underlying generation of this rhythm. We found the ketamine-related behavioral changes were attenuated by haloperidol, which has important translational value and suggests theta-driven HFO after ketamine, but not HFO generation *per se,* is dopamine-dependent. However, haloperidol can also act on NR2B receptors ^64^ and we cannot exclude the possibility that actions at this site contribute to the effect we observed.

Ketamine-dependent HFO correlated with stereotypic sniffing and hyperlocomotion, both widely established measures of psychotic-like behavior in rodents ^65,66^. Correlations between HFO power and hyperactivity have been reported by many groups, however a causal link has not been demonstrated. Interestingly, Hansen found that locomotion is not the major driving factor for ketamine-dependent HFO ^19^, in line with our previous work ^45^ (and haloperidol here) showing that reduced locomotor activity does not reduce HFO power. Here, we show that nasal respiration can drive the generation of ketamine-dependent HFO. This was demonstrated using simultaneous theremcouple and LFP recordings, and unilateral naris occlusion which was associated with unilateral reductions in HFO on the ipsilateral side. Ketamine-dependent faster stereotypic nasal respiration was associated with increased HFO power, but also slow basal respiration modulated HFO power in the presence of haloperidol. However, in the current study we did not examine the effect of low doses of ketamine, for example 3 mg/kg can produce weak increases in HFO without markedly altering locomotion ^20^. Further studies are warranted to document the effect of other psychoactive compounds such as LSD or DOI which can increase HFO power without affecting locomotion ^19,67^.

To date, the majority of studies examining NMDA receptor-dependent HFO and gamma rhythms have focused on their relevance to the NMDA hypofunction model of schizophrenia, which is supported by findings that both rhythms interact with antipsychotics and are changed in other models of schizophrenia. Several studies have shown gamma power increases in humans after ketamine, but frequencies above the gamma band are notoriously hard to record without invasive techniques. However, there is some evidence from MEG that ketamine-dependent HFO does occur in the human brain. Intriguingly, clinical reports have shown that the psychosis-like actions of ketamine correlate with later antidepressant effects ^6^ and can even predict clinical responses in treatment-resistant depression ^68^. The short-lasting changes in ketamine-dependent oscillations (around 15 min) may represent this initial phase of ketamine’s actions which lead to longer-term plastic changes.

HFO in the brain tends to denote spiking activity ^69^ and thus HFO synchronized at theta frequencies by respiration would be expected to reflect acute widespread firing of projection neurons. Indeed, ketamine-dependent HFO, at least in the OB, orchestrates the firing of projection neurons which appears essential for relay to limbic regions ^35^. Electrical high frequency stimulation is an emerging method used for the treatment of depression with effective frequencies commonly 130 Hz ^70,71^ strikingly similar to ketamine-dependent HFO. Against the notion that HFO reflect direct/indirect therapeutic effects, Zanos has shown that the antidepressant actions of the ketamine metabolite (2R,6R)-HNK are independent of NMDA receptor blockade and devoid of ketamine-related side effects ^72^. These data would argue that NMDA receptor-dependent actions of ketamine, including HFO, are not required for antidepressant effect, but rather may be associated with adverse drug effects. Thus although the functional relevance of ketamine-dependent oscillatory changes, whether therapeutic or adverse, remain to the explained the presence of fast oscillatory activity in the brain would be predicted to have physiological consequences. For example, *in vitro* electrical high frequency stimulation (HFS >100 Hz) has long been known to produce long-term potentiation (LTP) ^73^. Potentially more relevant to the ketamine model is that theta-burst high frequency stimulation also produces LTP ^74^, possibly in a more robust way ^75^, and appears necessary to evoke LTP in striatal tissue ^76^

It is worth noting that bulbectomy is a widely used rodent model of depression which produces downstream changes in the same limbic circuits as affected in patients ^36^. Ketamine-HFO are dependent on OB activity ^35^ and this issue gains important since intranasal delivery is the preferred route of administration for ketamine and its analogues, in depression. Thus, OB effects appear important which may work in parallel with other networks, for example Lang showed recently that ketamine inhibited burst activity in the habenula an “antireward” area ^71^. Although an OB-habenula pathway is believed to exist ^77^ it is unknown if these ketamine-dependent changes are related.

In summary, as noted by Moberly and colleagues, voluntary or involuntary changes in nasal breathing may potentially modulate brain activity ^26^. Olfactory sampling involves multiple brain regions and coherent oscillatory activity across areas may serve as a mechanism for temporal coordination of cortical and limbic networks ^36^. We conclude that nasal airflow is necessary for the emergence of ketamine-dependent HFO in multiple brain regions and behavioral hyperactivity. Zarate has proposed that mechanistic similarities may exist between ketamine-induced depersonalization and antidepressant responses ^68^. We speculate that ketamine’s HFO represent converging electrophysiological activity which could account for initial psychotic-like effects and later antidepressant effects and further studies are warranted to address this issue.

## Methods

Surgery: 25 male Wistar rats (250–350□g) were used in this study. Group 1 (thermocouple study): Twisted stainless steel electrodes (125□μm, Science Products, Germany) were implanted in the OB (AP□+7.5, ML□±0.5, DV 3–3.5□mm) along with two precision fine bare wire temperature sensors (80 μm diameter, 5TC-TT-KI-40-1M, Omega Engineering Inc., Czech Republic) into the right and left nasal cavity for monitoring nasal airflow (N=8). These rats were used to examine the effect of 20 mg/kg ketamine on nasal respiration and oscillatory activity in the OB. A subgroup (N=5 rats) were used to examine the effect of 1.0 mg/kg haloperidol pretreatment. Group 2 (multistructure study): Stainless steel electrodes were implanted in the OB, PFC (AP +3.2, ML 0.5, DV 3.0), and VS (AP 1.6, ML+1.0, DV 7.0) for simultaneous recording of LFPs used to examine HFO in multi-structures simultaneously (N=6). Two rats had electrodes bilaterally implanted in the OB, PFC, VS and were also used in naris blockade experiments (group 3 and 4). Four rats had additional electrodes targeted to the hippocampus and/or amygdala and piriform cortex for a supplementary experiment (ketamine-dependent HFO was inconsistent in these areas and the data not shown). Group 3 (unilateral naris blockade in PFC and VS): Five rats were implanted with twisted stainless steel electrodes bilaterally in the PFC and VS to determine the effect of unilateral naris blockade on these regions. Additionally, 2 rats from group 2 (bilaterally implanted in OB, PFC and VS) received naris blockade giving a total N=7. Group 4 (unilateral and bilateral naris blockade in OB): Stainless steel electrodes were implanted bilaterally in the OB of 6 rats for unilateral and bilateral naris occlusion experiments. Two rats from group 2 (bilateral implantation in OB, PFC, and VS) also received naris blockade giving a total N=8. A screw posterior to the bregma was used as a reference/ground in all cases.

One week after surgery, rats were placed in an arena (44×50×42 cm). LFPs and thermocouple recording were recorded through a JFET preamplifier, amplified ×1000, filtered 0.1–1000 Hz (A-M Systems, USA), digitized at 5 kHz (Micro1401, CED, Cambridge, UK). Horizontal locomotor activity was assessed by photocell beam breaks (Columbus Instruments, USA). Thermocouple experiments were performed according to the Latin-square design, whereby each rat was injected twice in a pseudorandomized order with either ketamine 20 mg/kg (Sigma, Poland) or saline. A subgroup of rats (N=5) were preinjected with 1 mg/kg haloperidol followed 15 min later by 20 mg/kg ketamine. Effect of 20 mg/kg ketamine on LFP oscillations was examined in rats with electrodes implanted in the OB, VS, PFC (N=6). For naris blockade experiments occlusion was achieved using a silicon occluder ^78^ either unilaterally for rats implanted with electrodes bilaterally in the VS and PFC (N=7). Rats with electrodes bilaterally in the OB received both unilateral and bilateral nares occlusion (OB; N=8). For naris blockade experiment, rats were baselined for 20 min and then briefly anesthetized using isoflurane to allow insertion of the silicon occluder(s). Rats were recorded for a further 60 min to ensure sufficient time for isoflurane washout, and then injected with ketamine 20 mg/kg. Electrode locations were determined on 40□μm Cresyl violet (Sigma, UK) or Hoechst (Sigma, UK) stained sections. All experiments were conducted in accordance with the European community guidelines on the Care and Use of Laboratory Animals (86/609/EEC) and approved by the 1^st^ Local Ethics Committee for Animal Experiments in Warsaw, Poland.

## Analysis

Mean power spectra of the LFP were computed on successive 30 s data blocks using a fast Fourier transform (4096 points) to calculate integrated, dominant power and frequency of low-gamma (30-60 Hz), high-gamma (70-100 Hz) and HFO (130–180□Hz). Thermocouple signals (1-10 Hz) were used to determine the dominant sniffing frequency and the proportion of fast (4-10 Hz) sniffing behavior. Waveform correlations of 130-180 Hz bandpass filtered LFPs ipsilateral (OB, PFC, VS) and contralateral OB were computed and maximum correlation and offset were used to calculate similarity and synchronicity.

### Phase-amplitude coupling analysis

Cross frequency phase-amplitude coupling was examined between the phase of oscillations in the thermocouple signal or low-frequency (1-10 Hz) LFP, and amplitude of high-frequency LFP oscillations (https://github.com/GabrielaJurkiewicz/ePAC). It enables analysis of coupling with the Modulation Index (MI) method ^79^ which relies on measuring the distance between the obtained distribution of high-frequency amplitude across low-frequency phases from the uniform one. The preprocessing of the LFPs and thermocouple data consisted of high-pass filtering (cutoff 0.1 Hz), lowpass filtering (cutoff frequency 250 Hz; both Butterworth 2nd order), downsampling to 625 Hz, and dividing data into 20 s fragments. The investigated low-frequency band ranged from 2-10 Hz (1 Hz step and 2 Hz filtration bandwidth) and high-frequency band ranged from 30-200 Hz (5 Hz step). Remaining parameters for the ePAC toolbox were set to default values (nbCycles=3, w=5, nbBins=18, Nboot=200, peMI=95, pPhaseCom=95, Athresh=0.1).

## Statistical analysis

Data are expressed as mean ± SEM or median ± interquartile range according to data normality. For multiple group analyses data were analyzed using repeated-measures ANOVA followed by the Bonferroni post hoc test or the Kruskal-Wallis test followed by Dunn’s multiple comparison test. Spearman’s rank correlation coefficient was used to examine the relationship between sniffing, locomotion and oscillatory activity. p<0.05 were considered statistically significant.

## Author contributions

Jacek Wrobel – Concept, acquisition, analysis, and interpretation of data

Władysław Średniawa - Analysis, and interpretation of data

Gabriela Bernatowicz - Analysis, and interpretation of data

Jaroslaw Zygierewicz - Analysis, and interpretation of data

Daniel K Wójcik - Analysis, and interpretation of data

Miles Adrian Whittington - Interpretation of data and drafting of manuscript

Mark Jeremy Hunt - Concept, acquisition, interpretation of data and drafting of manuscript

## Supplementary information

**Supplementary Figure S1. High-gamma and HFO lock to different phases of theta oscillations.** Example of the time course showing the effect of ketamine on the phase and strength of coupling of OB high-gamma (70-100 Hz, top) and HFO (130-180 Hz, middle) with respect to local theta. The bottom panel shows the mean and SEM for all rats. Injection of 20 mg/kg ketamine occurred around 20 min. Time courses show significant coupling only.

**Supplementary Figure S2. Unilateral naris occlusion does not affect ketamine-induced locomotion.** Time course of beam breaks after 20 mg/kg ketamine for mono-occluded naris rats and control non-occluded rats. Pre= pre-injection of ketamine.

